# Stable transgenesis in *Astyanax mexicanus* using the *Tol2* transposase system

**DOI:** 10.1101/535740

**Authors:** Bethany A. Stahl, Robert Peuß, Brittnee McDole, Alexander Kenzior, James B. Jaggard, Karin Gaudenz, Jaya Krishnan, Suzanne E. McGaugh, Erik R. Duboue, Alex C. Keene, Nicolas Rohner

## Abstract

*Astyanax mexicanus* is a well-established and widely used fish model system for evolutionary and developmental biology research. These fish exist as surface forms that inhabit rivers and 30 different populations of cavefish. The establishment of *A. mexicanus* as an emergent model organism for understanding the evolutionary basis of development and behavior has been accelerated by an increasing availability of genomic approaches to identify genotype-phenotype associations. Despite important progress in the deployment of new technologies, deep mechanistic insights into *A. mexicanus* evolution and development have been limited by a lack of transgenic lines commonly used in genetic model systems. Here, we expand the toolkit of transgenesis by characterizing two novel stable transgenic lines that were generated using the highly efficient *Tol2* system, commonly used to generate transgenic zebrafish. A stable transgenic line consisting of the zebrafish ubiquitin promoter fused to eGFP expressed eGFP ubiquitously throughout development in a surface population of *Astyanax*. To define specific cell-types, we injected fish with a Cntnap2-mCherry construct that labels lateral line mechanosensory neurons in zebrafish. Strikingly, both constructs appear to label the predicted cell types, suggesting many genetic tools and defined promoter regions in zebrafish are directly transferrable to cavefish. The lines provide proof-of-principle for the application of *Tol2* transgenic technology in *A. mexicanus*. Expansion on these initial transgenic lines will provide a platform to address broadly important problems in the quest to bridge the genotype to phenotype gap.

## Introduction

The Mexican tetra *Astyanax mexicanus* is a versatile model system, which is well suited for the study of the evolution of morphology, behavior, and physiology [1–10]. The different ecologies of the river and cave habitats of this species have led to the evolution of river dwelling and cave adapted populations which differ in a large number of traits (Figure 1a,b). The surface and cave morphs are diverged by ~200,000 years [11]. Despite the temporal and spatial divergence, the cave and surface populations remain interfertile. This ability may have been maintained by considerable gene flow between the surface and cave populations [11]. Many of the known cave populations are geographically isolated from each other and are the result of repeated evolution [11, 12]. Thus, one major question is whether the same genes or pathways are repeatedly modified throughout evolution. Importantly, the fish can be kept in the laboratory and breed regularly. As such they are likely amenable to most procedures that are available for other fish model species, such as the zebrafish (*Danio rerio*) or the medaka (*Oryzias latipes*).

**Figure 1.**
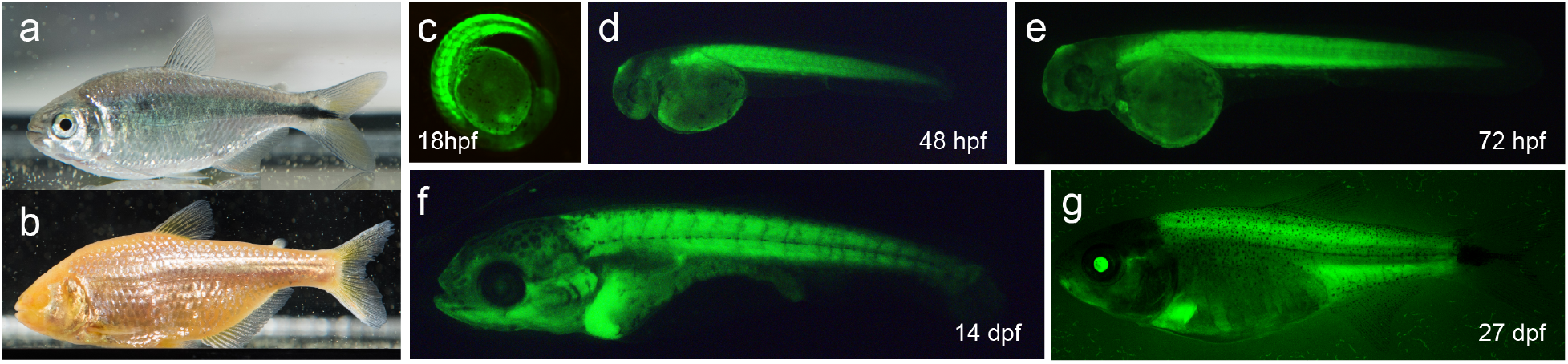
Ubi-GFP stable transgenic line expresses GFP at different developmental stages of *Astyanax mexicanus*. a, b) Surface fish (a) and cavefish (b) forms of *Astyanax mexicanus* are a versatile model system for evolutionary and developmental biology research. c-g) GFP expression in different developmental stages of surface fish of transgenic *A. mexicanus*. c) 18hpf d) 48hpf e) 72hpf f) 14dpf g) 27 dpf. hpf = hours post fertilization, dpf = days post fertilization (images not to scale).

Over the last decade, an array of genomic and genetic tools have been established, which allow for mechanistic investigation into the differences between surface fish and cavefish. High-resolution genetic maps [13–15], data from transcriptome analyses [16, 17], methods for altering gene expression [18], and approaches for introducing mutations [19, 20] have recently been applied to study the genetic basis of trait evolution. Moreover, the genomes of multiple *A. mexicanus* populations have recently been assembled, further developing *Astyanax mexicanus* as a tractable model system [11, 21]. Despite these technological advances, the system suffers from the limited availability of transgenic lines. To date, only one stable transgenic line has been published that was generated using the I-SceI-meganuclease system [22]. A few studies have used *Tol2* technology, commonly used to generate transgenic zebrafish, but analysis was limited to mosaic F0 fish [6, 22], therefore, procedures for generating stable transgenic lines are unclear.

Multiple features suggest *A. mexicanus* are well-suited for the implementation of high throughput transgenic technology. These fish have large clutch sizes of externally fertilized eggs and a short generation time of 5-6 months [23]. Further, *A. mexicanus* are closely related to zebrafish, with an estimated divergence of 100 million years [24]. A large array of transgenic lines from zebrafish are available that depends on *Tol2* mediated transgenesis. If the same *Tol2* transgenic constructs that are used in zebrafish could be used for *Astyanax*, this could create important synergies. Here, we report the successful generation of two new stable transgenic lines in *Astyanax mexicanus*. Both lines took advantage of zebrafish promoters; one line drives *eGFP* under a ubiquitous promoter and one line drives mCherry in lateral line neuromasts and neurons within the brain. Both lines resemble the expression pattern and timing of the zebrafish lines, strengthening the possibility of cross-species applicability.

## Results

Promoters that drive transgene expression ubiquitously provide important tools for every model organism because it allows for cell fate and lineage tracing experiments [25]. To optimize *Tol2* transgenesis, we first injected fish with the *Tol2* construct that included 3.5 kb of the zebrafish ubiquitin promoter upstream of eGFP (Ubi-GFP) [26]. This construct was co-injected with *Tol2* mRNA to *A. mexicanus* surface fish eggs ages 0-60 minutes post fertilization. We noted broad mosaic expression in many F0 fish, providing an indicator of successful construct injection. These mosaic fish were selected for breeding as adults to identify stable transgenic insertions.

Pair-breeding revealed that female F0 fish produced 10% GFP-positive progeny (F1) (n=65/650). We raised the F1 fish to maturity and in-crossed individual fish to generate the F2 generation. These crosses produced an average of approximately 75% GFP positive F2 embryos, in line with both carriers being heterozygous for the transgene. To confirm that the zebrafish promoter is ubiquitously active, similar as in zebrafish, we checked different developmental stages for GFP expression. We detected ubiquitous expression in all developmental stages tested from 18 hours post fertilization (hpf) to 27 days post fertilization (dpf) (Figure 1c-g).

To test if the ubiquitin promoter is active in different tissues we sectioned transgenic larvae at 14 dpf and measured GFP expression in transverse sections. While we detect slight differences in the intensity of GFP expression in sections, there is overall ubiquitous expression throughout the embryo. We next dissected brain, liver, eyes, and fins of an adult fish and all tissues displayed strong expression of GFP (Figure 2). Figure 2a displays a transverse section at the level of the eyes. One important advantage of the zebrafish Ubi-GFP line, distinguishing it from other available ubiquitous lines, is that the promoter is active in all major blood cell types [26]. To test if this is true for *A. mexicanus*, we dissected the haemopoietic organ, the head kidney, of an adult transgenic fish and measured expression of GFP in hematopoietic cells using a flow cytometry approach. All major cell populations that are distinguishable by forward scatter and side scatter (erythrocytes, myeolomonocytes, lymphocytes, and precursors) display strong GFP expression, similar to what has been observed in zebrafish [26] (Figure 2c). This broad multilineage expression makes the ubi-GFP line a suitable reagent for hematopoietic transplantation experiments.

**Figure 2.**
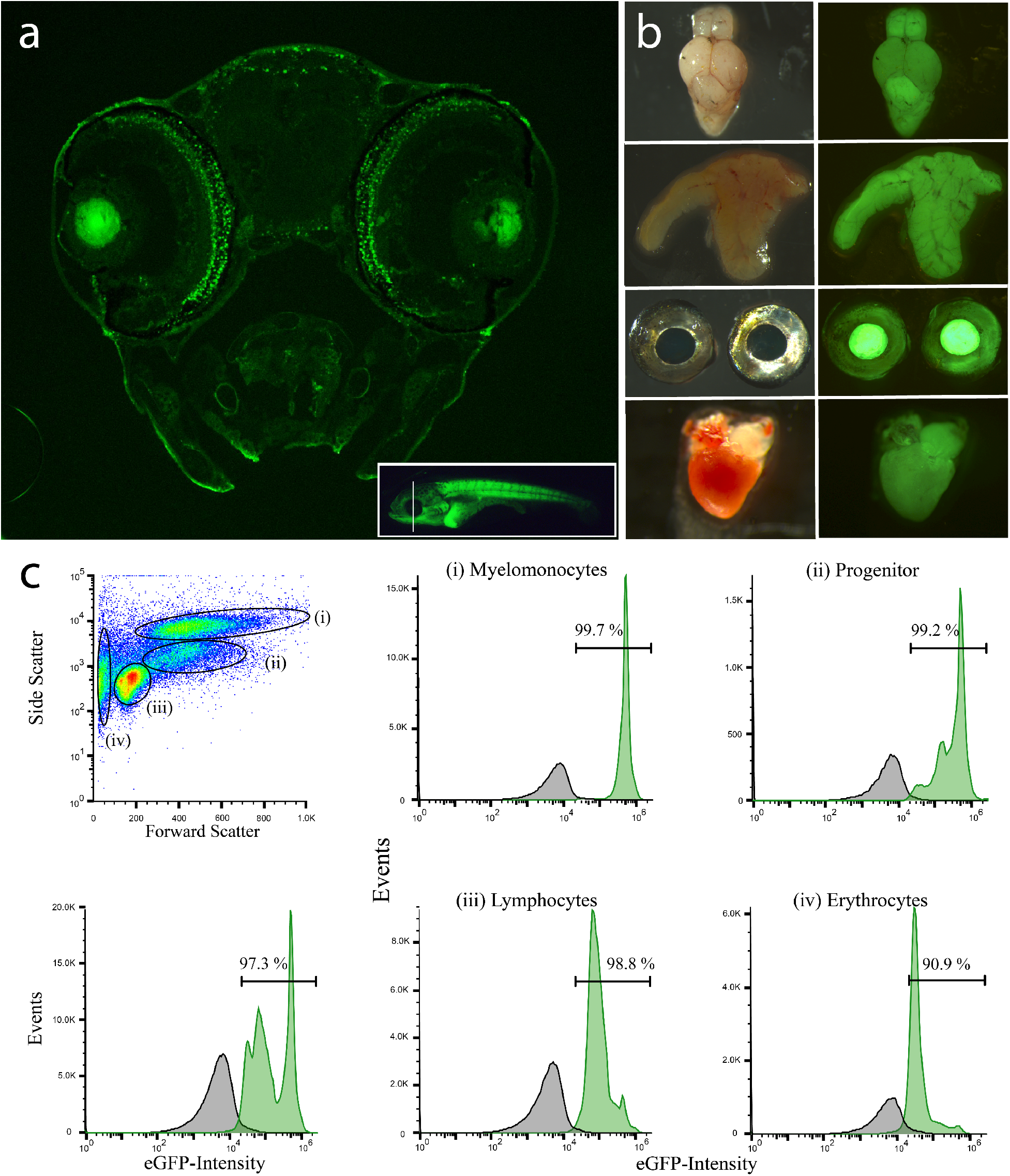
Ubiquitin promoter driven transgene expresses GFP in different tissues. a) transverse section of a 14dpf old Ubi-GFP transgenic surface fish. Inset picture indicates the location of the section. b) GFP expression in different organs of adult transgenic fish, from top to bottom: brain, liver, eyes, heart. c) Fluorescence activated cell sorting of head kidney cells from Ubi-GFP transgenic *A. mexicanus*. Four distinct hematopoietic cell populations can be identified by forward scatter and side scatter: (i) myelomonocytes, (ii) progenitors, (iii) lymphocytes and (iv) erythrocytes. Green indicates GFP signal from Ubi-GFP transgenes, while dark grey indicates GFP signal from wild-type control.

A common application of transgenesis is the targeting of reporters to specified tissue using tissue-specific promoters. In *A. mexicanus*, the lateral line is thought to underlie the evolution of many behaviors including foraging, social behavior, and sleep [27]. In zebrafish, a transgenic construct consisting of a 2Kb promoter fragment of contactin associated protein like 2a (Cntnap2a) driving mCherry (Cntnap2a:mCherry) labels the lateral line afferent neurons at 3 dpf, but expression patterns later in development have not been reported.

We injected *A. mexicanus* surface fish with the zebrafish Cntnap2a:mCherry promoter, isolated mosaic F0 fish, and screened for stable insertions. We found modest labeling of the lateral line and multiple brain regions including the mesencephalon, and rhombencephalon as early as 2 days post fertilization (dpf), with expression becoming more prominent after 7 dpf (Fig 3a-i). At all time points, lateral line neuromasts that project into the brain can be clearly detected and persist through 20dpf fish (Fig 3m-r). The signal in the gut is likely autofluorescence, as it was visible in non-injected fish (data not shown), and no other visible signal was detected outside of the central nervous system. Across developmental time points, Cntnap2a:mCherry selectively labeled portions of the brain including the hindbrain regions such as the medial octavolateral nucleus, that is known to receive inputs from the lateral line in other fish species [28, 29]. In addition, labeling was detected in the optic tectum and olfactory bulb after 5dpf, suggestion the expression pattern is not limited to the lateral line.

**Figure 3.**
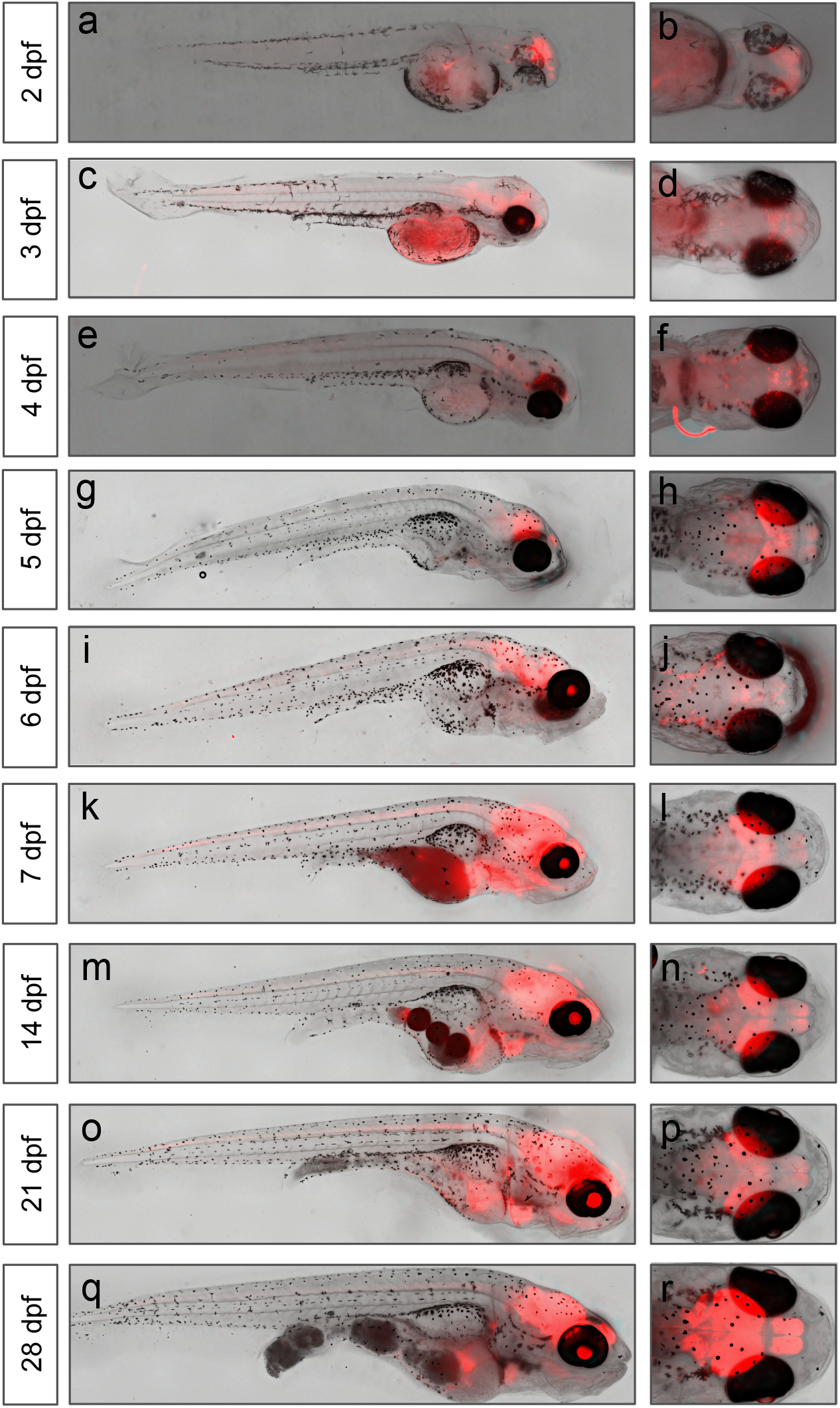
Expression of Cntnap2a-mCherry across development. Visualization of Cntnap2a-mCherry expression during early development (2 dpf – 28 dpf) in whole body (left column) and dorsal head view (right column). Cntnap2a expression is evident throughout the lateral line and hindbrain regions.

To verify that Cntnap2a-mCherry labels the lateral line, we imaged 6 dpf Cntnap2a-mCherry transgenic fish co-stained with DASPEI, a dye that labels mitochondria and is widely used to mark lateral line neuromasts in diverse fish species (Fig 4a). Colocalization between DASPEI and mCherry label was detected in neuromasts along the body (Figure 4b). In addition, close-up imaging reveals projections from multiple neuromasts forming a lateral line nerve that innervates the brain (Figure 4d). To determine the expression pattern of Cntnap2a-mCherry within the brain, whole mount confocal images of 6dpf fish co-labeled with tERK were acquired. Anatomical segmentations were performed to localize the expression pattern of Cntnap2a-mCherry. This revealed strong expression of hindbrain areas in the rhombencephalon such as the medial octavolateralis nucleus (MON) and several other areas likely associated with sensory processing in the hindbrain. There was additional strong mCherry expression within regions of the mesencephalon and telencephalon with further sparse labeling within the diencephalon. Together, these findings verify that Cntnap2a-mCherry transgene labels lateral line sensory neuromasts and projects centrally into several distinct nuclei, providing proof-of-principle for selective expression using a zebrafish promoter-fusion construct in *A. mexicanus*.

**Figure 4.**
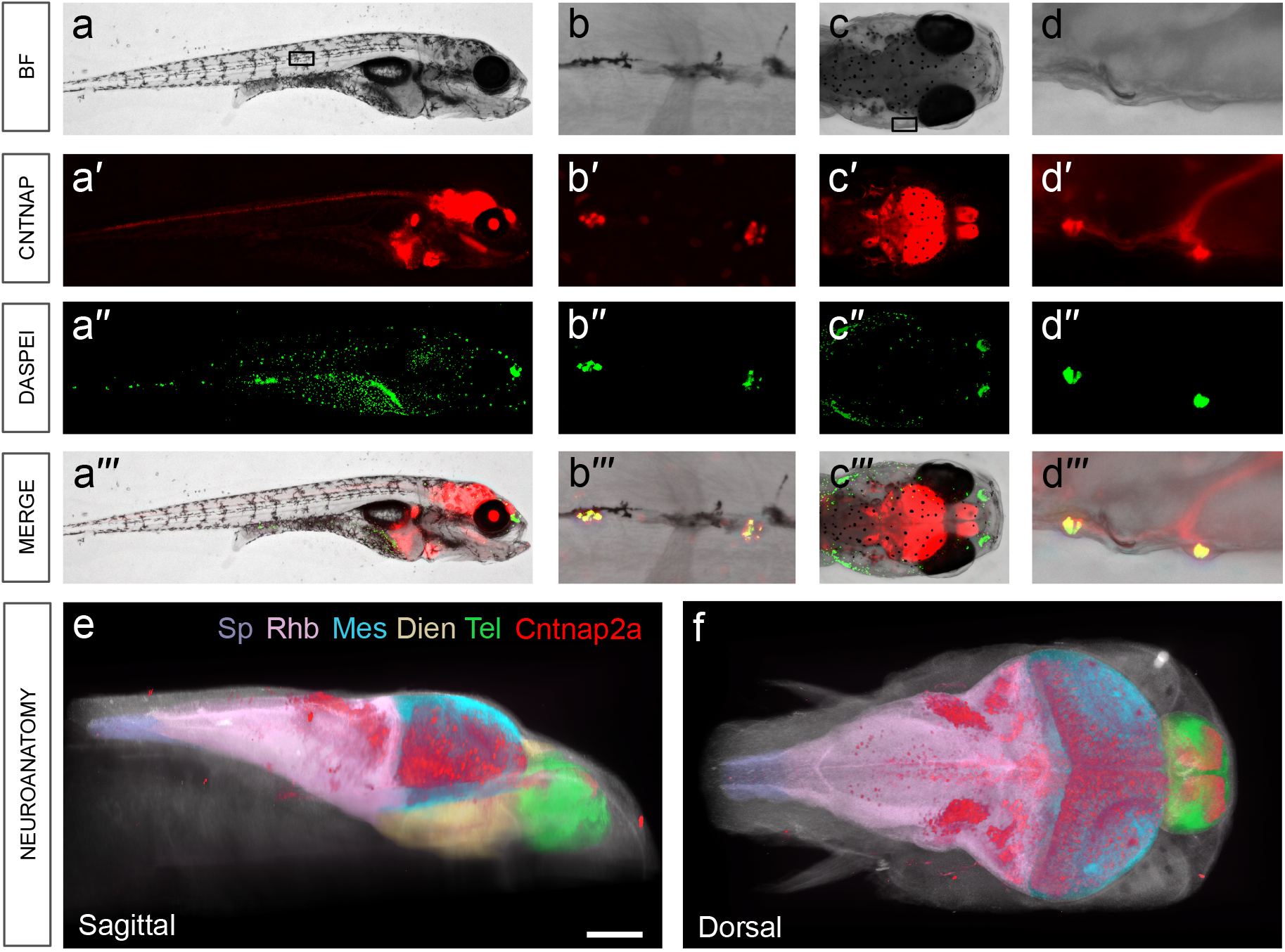
Cntnap2a colocalizes with DASPEI neuromast stain. a) Whole body view and b) zoom shows that Cntnap2a-mCherry (red) and neuromasts of lateral line (DASPEI, green) colocalize (merge, b’”). c) Dorsal head view and d) zoom demonstrates colocalization of neuromasts and Cntnap2a labeled neurons that innervate the brain (merge, d’’’). e) Sagittal aspect. f) Dorsal aspect (SB= 100 μm), Spine (Sp, purple), Rhombencephalon (Rbh, pink), Mesencephalon (Mes, blue), Diencephalon (Dien, yellow), Telencephalon (Tel, green), Cntnap2a:mCherry (red).

## Discussion

Here we describe two new transgenic *Astyanax* lines, one that ubiquitously expresses GFP in all tested developmental stages and tissues and one that is more selective. In the latter, mCherry expression is limited to the central nervous system, labeling the lateral line and other selective regions within the brain.

The use of transgenic lines is important to label tissues and cell types of interest and to overexpress allelic variants under the control of specific promoters or enhancers. *Astyanax mexicanus* is well suited for the generation of transgenic lines: the fish produce large clutch sizes of externally fertilized eggs with a penetrable chorion that permits easy microinjection. Breeding occurs frequently and regularly under lab conditions (usually every 1-2 weeks). There are protocols for *in vitro fertilization* that permit the generation of timed embryos and the fish have a relatively short generation time [23, 30]. To our knowledge, this is the first study that uses *Tol2* for the generation of stable transgenic lines in *Astyanax mexicanus. Tol2* possesses important advantages over alternative transgenic methods and strongly increases the insertion efficiency of the recombinant DNA and germline transmission [31]. Of particular relevance is that we did not detect significant differences in expression between the zebrafish and the *Astyanax* lines minimizing concerns about the evolutionary divergence of promoters between the two species. Currently, there are over 40,000 transgenic lines described in zebrafish (ZFIN) which opens vast possibilities for *Astyanax* and EvoDevo research.

The Ubi-GFP line can be used for cell fate and lineage tracing experiments or to test the autonomy of cell and tissue signaling on certain phenotypes. For example, the line can be used to transplant cells from surface fish to cavefish and monitor the behavior of the clones [32]. While we did not detect obvious differences in expression levels in different tissues, a ubiquitous line does not guarantee homogenous expression. In zebrafish, for example, it has been shown that the construct is expressed lower in certain subsets of internal organs such as some subsets of neurons and epithelial cells of kidney tubules [26]. Similarly, we observed slight variability in the intensity of GPF expression in different tissues and areas of the embryos.

Additionally, we find that the zebrafish Cntnap2a:mCherry promoter labels the lateral line in *A. mexicanus*. Unlike many other behaviorally relevant neuronal cell populations in *A. mexicanus*, the lateral line has been well studied because it can be labeled using the mitochondrial cilia marker DASPEI, and neuromasts can be ablated with the ototoxic antibiotic gentamicin. These approaches have been used to characterize functional differences of the lateral line between surface and cavefish. An expansion of the lateral line is thought to underlie the reduced dependence on visual processing in cavefish, and confer increased vibration sensitivity and reduced sleep, presumably to improve foraging [33–35]. Genetic labeling of the lateral line using the transgenic fish is highly superior to physical labeling with DASPEI because it allows more fine-scale assessment of neurons involved in lateral line function between cavefish and surface fish. The lateral line neuromast projections can be examined for anatomical differences between populations of *A. mexicanus*. In addition, the Cntnap2a promoter driving the Ca^2+^ sensor GCaMP would allow for Ca^2+^ to functionally compare lateral line activity.

In zebrafish, significant headway has been made identifying neural circuits that regulate behavior. The identification of these circuits allows for cell-autonomous manipulation of gene function, or optogenetic manipulation of behaviorally relevant neurons. In *A. mexicanus* a number of genes have been implicated in behavior including a role for Mc4r in feeding, hypocretin in sleep, and monoaminergic signaling in aggression [6, 36, 37]. While presumably the neurons expressing these genes are implicated in behavior, a lack of transgenic accessibility represents a fundamental impediment to mechanistic investigation. The proof-of-principle of efficient implementation of *Tol2* transgenesis in *A. mexicanus* should allow for the implementation of binary transgenic expression systems commonly used in other genetic model systems including GAL4/UAS and QF/qUAS systems [38, 39]. In addition to promoter fusions, these systems would allow for developing libraries of transgenic effector lines to manipulate neuronal function. The transparency, generation time, and standardized husbandry procedures currently available in *A. mexicanus* strongly position this emergent model system for the implementation of these approaches.

The generation of transgenesis and gene-editing methodology is growing increasingly common in fish species other than classical models like zebrafish and medaka. For example, in the African cichlid, *A. burtoni* [40], the Midas cichlid, *A. citrinellus* [41], the short-lived African killifish *N. furzeri* [42] or the tilapia *O. niloticus* [43] transgenesis has been reported. Nevertheless, the application of these techniques is limited to a few lines, while the power of genetic approaches in model organisms typically derives from the generation of transgenic libraries consisting of thousands of lines. While the optimization of methodology provides a platform for developing these lines, the success is also dependent on community resources such as stock centers and large-scale initiatives to generate lines. Our finding that constructs are directly transferable from zebrafish to *A. mexicanus* may streamline the process of generating these lines.

## Acknowledgements

We thank Christian Mosimann and Jenna Galloway for the Ubi-GFP construct and Hernan Lopez-Schier for the Cntnap2a-mCherry (SILL1) construct. We thank the Aquatics facility of the Stowers Institute for care and maintenance of the fish, in particular Zachary Zakibe for help with IVF and Alba Aparicio for counting embryos. We thank the Histology and Microscopy facility at the Stowers Institute for assistance with the data acquisition. Original data underlying this manuscript can be accessed from the Stowers Original Data Repository at http://www.stowers.org/research/publications/libpb-1381. Funding from the Stowers Institute and the Edward Mallinckrodt foundation to NR, NIH R21 NS105071 to ACK and ERD, NIH R01 GM1278872 to ACK, SEM and NR and NSF 1656574 to ACK supported this work. R.P. was supported by a grant from the Deutsche Forschungsgemeinschaft (PE 2807/1-1).

## Author contributions

NR, AK, ED, SM conceived and designed the analysis. BS, RP, BM, AK, JJ, KG, JK, performed the experiments. BS, AK, NR wrote the paper with input from all other authors.

## Methods

### Fish husbandry and egg collection

For *Astyanax* F0 transgenesis, fish are bred in community breeding tanks consisting of 2-3 females and 3-5 males. To promote breeding, fish are fed a high fat diet starting ~1 week prior to injections and heaters with +2° C temperature increase are added 2 days before breeding night, in accordance with previously published methods [44]. On the night of breeding, tanks are frequently examined for fresh, single cell eggs. Eggs are collected with a fine mesh net and loaded on to injection plates. Injection plates are prepared using 3% agarose dissolved in 100 ml fish water, poured into a 100 ml petri dish, and an egg injection mold (Adaptive Science Tools) is added to create wells for the eggs.

### Constructs

#### Cntnap2a-mCherry

We utilized the previously published zebrafish mCherry labeled Cntnap2a construct Tg(hsp70l:mCherry-2.0cntnap2a); zfin #ZDB-TGCONSTRCT-120424-6. In zebrafish, placing a 2kb Cntnap2a enhancer fragment downstream of the heat shock-70 promoter and mCherry drove robust expression in lateral afferent neurons. This enhancer fragment is also known as the sensory innervation of the lateral line number 1 (SILL1) promoter [45].

#### Ubi-eGFP

We utilized the previously published zebrafish eGFP labeled ubiquitin promoter construct Tg(−3.5ubb:EGFP); zfin ZDB-TGCONSTRCT-110317-1. This construct contains a 3.5 kb promoter fragment upstream of the ATG of the zebrafish gene ubiquitin B (*ubi*), including the first non-coding exon and the intron. It has been shown to ubiquitously express eGFP in most tissues and developmental stages including the hematopoietic organ [26].

### RNA synthesis of *tol2* transposase RNA

The *Tol2* transposase plasmid (pCS-zT2TP) was used as a template for *Tol2* mRNA generation. The pCS-zT2P plasmid was digested with restriction enzyme and purified. *In vitro* transcription of *Tol2* mRNA and lithium chloride precipitation/purification was performed using the SP6 mMessenger Kit (Ambion #AM1340) according to manufacturers guidelines.

### Microinjection

Pull needles for injection using borosilicate glass capillaries (Sutter Instruments #BF100-58-10) in an electrode/needle puller (Sutter Instruments Model P-97). This protocol will vary per machine, but a sample needle-pulling program is: Heat 510, Pull 55, Velocity 100, Time 40, Pressure 500, Ramp 534. Back fill needles with approximately 3-4 ul of *Tol2* mRNA and the desired plasmid at a concentration of 25 ng/ul each in RNase-free water. Filled needles are loaded in a micromanipulator (World Precision Instruments #M3301R) connected to a microinjector/picoinjector (Warner Instruments #PLI-100A). Injection pressure settings were optimized for injections in a 1 nl volume and eggs are injected at the single cell stage.

### DASPEI stain

6 dpf Cntnap2a-positive fish were treated for 1 hour with 0.05% DASPEI (2-4-dimethylamino-N-ethylpyridinium iodide; Sigma Aldrich), which labels both superficial and canal neuromasts [46]. The precise mechanism of DASPEI labeling remains unknown, it is proposed that the dye enters the cells through transduction channels and apical endocytosis, allowing for uptake by active hair cells, making it specific for labeling intact neuromasts of the lateral line [46].

### Live Imaging

Live cavefish was used to image ubiquitin-GFP transgenic fish. All fish were anesthetized using MS-222 (Tricaine). Embryos were mounted in 3% hydroxypropyl methyl cellulose (VWR, cat# 200012-722) in E3 medium (ref). Juveniles were placed on a filter paper on top of a MS-222 soaked sponge. Images were acquired using a Leica M205C stereo microscope (Leica Microsystems Inc.) equipped with a filter set for detection of GFP (WB). The zoom setting for embryos was 4.0 and for juveniles 1.25. Images were processed using the LAS AF ver. 4.7 software (Leica Microsystems Inc.) and ImageJ.

### Fixed Imaging

*Astyanax* surface F1 fish expressing Cntnap2a-mCherry at various developmental time points were fixed overnight in 4% paraformaldehyde and rinsed in PBS solution. Fish were mounted in 2% low melt agarose and imaged at 10x or 20x magnification with a Keyence BZ-X710 invert microscope. Images were acquired with BZ Capture and processed using BZ-X analyzer software. Fish co-labelled with tERK antibody (###) were treated with 150 mM Tris-HCl at 70°C for 15 minutes, incubated in 0.05% Trypsin-EDTA for 35 minutes on ice. Fish were then incubated in primary antibody overnight at 4°C and for 95 minutes in secondary antibody at room temperature. Fish were then imaged on a Nikon A1 confocal microscope. All anatomical segmentations were performed using Amira 6.2 and visualized with a Volren module.

### Preparation of cryo sections for GFP imaging

14 dpf fish embryos were embedded with OCT compound (cat# 4583, Tissue-Tek, CA) and snap-frozen at −80C. Sections are prepared as previously reported [47]. Briefly, cryo sections with 12um-in-thickness were cut using a Cryostar Nx70 Cryostat (cat# 957020, Thermo Fisher Scientific) and picked onto superfrost microscope slides (cat# 12-550-15, Thermo Fisher Scientific). After 2 hours drying slides are immersed into PBS to wash off OCT, followed by coverslipping with ProLong gold antifade mountant (cat# P36930, Thermo Fisher Scientific). GFP imaging of slides and fish organs was performed with Axioplan2 (Zeiss) and Leica M205C microscopes.

### Hematopoietic tissue analysis

One 1 year old Ubi-GFP transgene fish and one similar aged wild type fish were euthanized with cold 500 mg/L MS-222 for 10 min. Hematopoietic tissue (head kidney) was dissected and tissue was forced through 40 μm cell strainer in L-15 media containing 5mM HEPES (pH 7.2), 10 % sterile water and 20 U/mL Heparin. Cell strainer was washed once, and single cell solution was washed once and resuspended. Flow cytometry analysis was based on forward scatter and side scatter and FL1 (GFP) with an EC-800 flow cytometer (Sony Biotechnology).

